# Self-organization of Computation in Neural Systems

**DOI:** 10.1101/016725

**Authors:** Christian Tetzlaff, Sakyasingha Dasgupta, Tomas Kulvicius, Florentin Wörgötter

**Affiliations:** Institute for Physics – Biophysics, Georg-August-University, Friedrich-Hund Platz 1, 37077, Göttingen, Germany; Bernstein Center for Computational Neuroscience, Georg-August-University, Friedrich-Hund Platz 1, 37077, Göttingen, Germany

## Abstract

When learning a complex task our nervous system self-organizes large groups of neurons into coherent dynamic activity patterns. During this, a cell assembly network with multiple, simultaneously active, and computationally powerful assemblies is formed; a process which is so far not understood. Here we show that the combination of synaptic plasticity with the slower process of synaptic scaling achieves formation of such assembly networks. This type of self-organization allows executing a difficult, six degrees of freedom, manipulation task with a robot where assemblies need to learn computing complex non-linear transforms and – for execution – must cooperate with each other without interference. This mechanism, thus, permits for the first time the guided self-organization of computationally powerful sub-structures in dynamic networks for behavior control.

---

When we are performing a complex skill, like neatly stacking two blocks, our motor system needs to accurately control position and orientation of the hand, which is a process that took us quite some time to learn when we were children. During learning, synaptic plasticity in the nervous system forms functional networks – often called ”cell assemblies” – that allow us to perform motor fine-control. Several thousands of neurons in many cell assemblies are active during any motor task and perform complex non-linear calculations to control the different degrees of freedom of the respective limbs. Adults master a large number of motor skills requiring a multitude of different cell assemblies most – if not all – of which have been formed by learning. To achieve such mastery, our brain has to solve a very complex problem. It needs to create a large number of computationally very powerful assemblies, which can only be achieved by using relatively small quantity of neurons for any one of them and by involving the same neurons in many different motor tasks. How such interwoven assembly networks are formed and how powerful assemblies can coexist therein without catastrophically interfering with each other remains unknown. Understanding this would, thus, carry substantial promise for our comprehension of how the brain can self-organize and provide the required requisite variety for complex motor control (*1*).

It is known, on the one hand, that networks can be trained to perform complex nonlinear calculations (*2, 3*), which could be used for motor control. This requires that those networks produce a reservoir of rich, transient dynamics from which the desired outputs can be siphoned off (*4*). On the other hand, it is also easy to create cell assemblies using hebbian learning rules that strengthen a synapse if preand post-synaptic neurons are co-active within a small enough time window (*5, 6*). It appears straight-forward to combine those mechanisms to arrive at the required assembly networks. Alas, two effects can destroy such an approach. Self-organization of neurons into cell assemblies by the processes of synaptic plasticity induces ordered or even synchronized neuronal dynamics replicating basic processes of long-term memory (*7–9*). This will reduce the dynamics of the network often to a degree that the required requisite variety for complex calculations cannot be provided by it any longer (*10*). In addition, trying to simultaneously create multiple assemblies will lead indeed to the aforementioned catastrophic interference if one cannot prevent them from growing into each other.

In this study, we exploit for the first time the interaction between neuronal and synaptic processes acting on different time scales to enable, on a long time scale, the self-organized formation of assembly networks, while on a shorter time scale, to conjointly perform several non-linear calculations needed to control six degrees of freedom of a motor system (robot) in parallel.

To understand how this can be achieved, we first show that assembly growth will lead to improved computational power using an example where a growing assembly is required to calculate some freely chosen non-linear transforms (e.g., power of 7 of the input). Crucial for this is that assembly growths must not create persistent or even synchronized network activity by which the required requisite variety of the assembly dynamics would be destroyed.

An external signal repeatedly stimulates several randomly chosen rate-coded input neurons in a randomly and very weakly excitatorily connected neuronal circuit with dominating inhibition (Figure 1 A and Materials and Methods). Every stimulation induces synaptic strengthening and thereby the outgrowth of a cell assembly starting from the input neurons (Figure 1 B and supporting online material [SOM]). Synaptic weights and – as a consequence – neural activities remain limited by the interaction of hebbian synaptic plasticity (long-term potentiation [LTP]; (*12*)) with the slower mechanism of synaptic scaling (*13, 14*). This aspect will be of direct relevance when later on considering several assemblies in parallel. Importantly, due to the dominating inhibition, the here formed cell assembly does not serve as an attractor of the neuronal dynamics and it does not produce persistent neuronal activities (*9*), which would be detrimental to computation (see SOM). Thus, with this type of repeated stimulation we obtain a growing assembly with transient activity.

**Figure 1:**
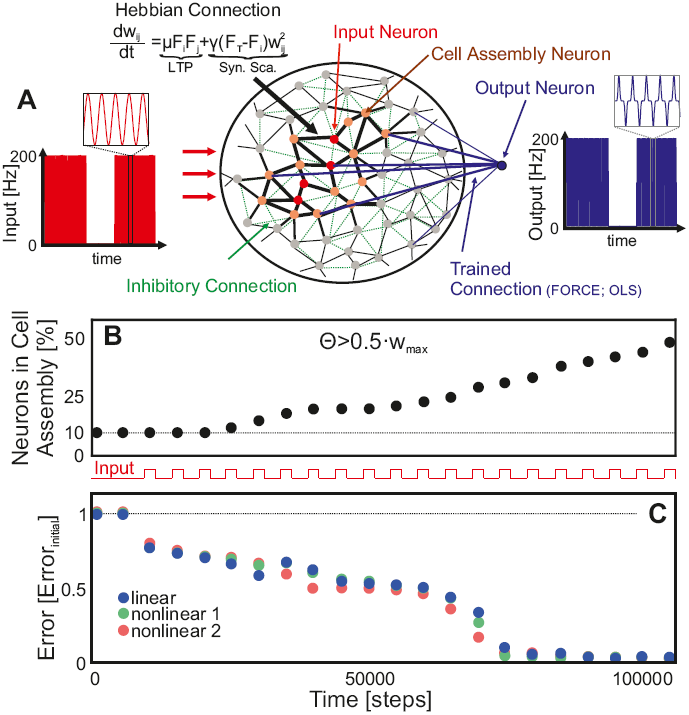
Cell assembly size and computational performance are correlated. (A) An input is delivered to several neurons (red disks) in a random neuronal network with very weak but plastic excitatory (black lines) and constant inhibitory connections (green dashed lines). The interaction of long-term potentiation (LTP) and synaptic scaling (Syn. Sca.) enables the formation of a cell assembly (red and orange disks) by increasing synaptic efficacies (thicker lines) when repeating the input several times. Note this network is not topographically organized. The here shown neighborhood ordering is for graphical reasons only. Output neurons (blue disk) are connected (blue dashed) to the full network and trained in a conventional way to create the desired output (by FORCE (*11*) or ordinary least squares (OLS) (*2*)). Here we used a single output neuron, but several can be connected without additional constraints (see also Fig. 3). (B) With more learning trials the assembly grows and integrates more neurons. To measure this, we arbitrarily define assembly size by that set of neurons connected with efficacies larger than Θ = 0.5·*W*_*max*_. (C) Parallel to the outgrowth of the cell assembly the error of the system to perform several linear and non-linear calculations decreases. linear: *O*(*t*) = *I*(*t*); nonlinear 1: *O*(*t*) = *I*(*t*)^3^; nonlinear 2: *O*(*t*) = *I*(*t*)^7^; *O*(*t*): output; *I*(*t*): input; please see SOM for protocol details.

Now we can analyse the computational power of such a structure during the growth process. To this end we interrupted assembly growth after each stimulus presentation and trained an output neuron connected to the *whole* network to perform one of several linear or non-linear calculations (*2, 3, 11*) and measured the error. We observed that the performance of the network strongly correlates with the size of the cell assembly (compare Figure 1 B to C).

Comparing the self-organized assembly network to networks with unchanging synapses (”static” networks) shows that it is indeed the embedding of a strongly connected assembly that creates the computational power. Computational reservoirs are commonly generated using random connectivity of variable strength between many neurons (*2, 3*) and, as a result, weak and strong synapses are randomly distributed therein. Inputs are also provided to randomly chosen units (*2, 3*). Consequentially, an input will lead to a spatially extended activity trace including many neurons if and only if one makes sure that there is a large-enough number of strong synapses existing in the network to begin with. In these cases the error, when performing a computation, is indeed low. The data highlighted by a light blue box in Figure 2 A shows that randomly connected static networks with few strong synapses (Fig. 2 B) will perform poorly as compared to networks with many strong synapses (dark blue box Fig. 2 A and Fig. 2 C), where the error is essentially zero. Remarkably, the same small error is obtained with an assembly network after some growth (Fig. 2 D) with only a small fraction of strong synapses, which can be seen when comparing the yellow (early learning) with the orange box (late learning) in Figure 2 A. The inputs provided to the assembly penetrate deep because strong synapses exist mainly within the assembly and this creates the required rich dynamics. This is beneficial for several reasons. Limiting the number of randomly distributed strong synapses mitigates the problem of potential run-away activity (*15*), which would entirely destroy computation. In addition, possibly the most interesting property of such assembly networks is that, due to the limited number of strong synapses per assembly, several assemblies can coexist and/or compete in a realistic way with each other. As shown above, synaptic scaling counterbalances the unrestricted growth processes of hebbian learning guaranteeing that the system stays in the non-persistent activity regime (*9, 15*). But, in addition, this also creates a competition between different presynaptic sites (*16*). This competition, which takes place on a neuronal scale, leads to a competition/coexistence between cell assemblies on the network scale. Hence, with an alternating, balanced presentation of two stimuli (”A”,”B”) two assemblies embedded in the same network grow in the same way (Fig. 2 F before trial 50), but with dominant presentation of stimulus ”A” the corresponding assembly will take over (Fig. 2 F after trial 50) without interference between input traces. This is difficult to achieve in a random network as one has no control over the actual input trace configurations, which might easily randomly interfere and perturb each other (*10, 17*).

**Figure 2:**
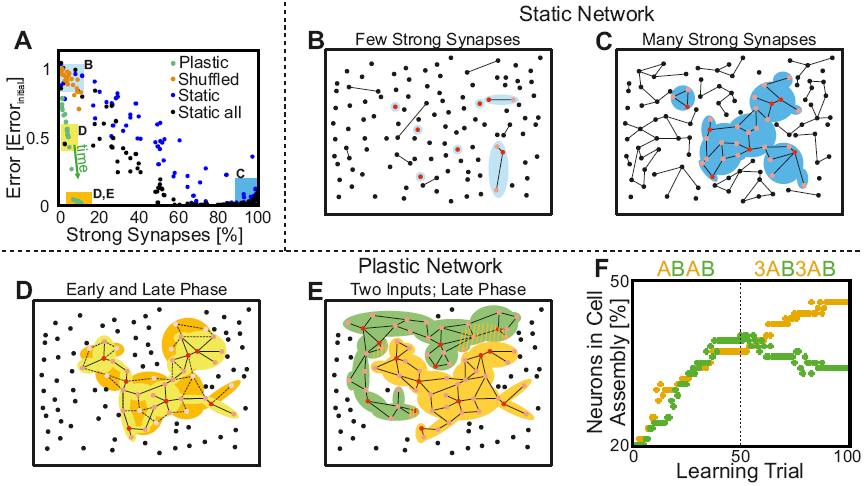
Comparison of the computational power of different networks relative to their synaptic structure. (A) The error in performing non-linear calculations (here nonlinear 1) decreases with the number of strong synapses having a weight of *W* > 0.5·*W*_*max*_ in the network. We created 300 randomly connected but unchanging networks (”static”) with 100 neurons each; plotting the error they produce against their number of strong synapses (blue, black dots). Green dots show the error from networks that are obtained during the temporal development of assembly formation (temporal progress indicated by the arrow). The assembly network (green) needs far fewer strong synapses as compared to the randomly connected static structures to achieve small errors (black: all neurons receive input; blue: same neurons as in the assembly network receive input). Shuffling the weights in the assembly network (orange dots) leads to the same low performance as for the static networks demonstrating that random arrangements of the same strong synapses does not suffice. (B-E) Schematic illustration of the underlying topology for different networks (red dots: stimulated neurons; orange dots: non-stimulated neurons driven by the input due to strong enough synapses; black lines: strong synapses; colored shadings: regions driven by the input). (B) static network with few strong synapses, (C) static network with many strong synapses, (D) plastic network after few learning trials (yellow shading) and after many learning trials (yellow+orange shading; dashed lines=strong synapses obtained by longer learning). The color coded boxes in panel (A) show the errors for cases (B-D). (E) Schematic of a plastic network with two cell assemblies competing for neurons (striped areas). (F) Competitive development of the two competing cell assemblies ”A” and ”B” as a function of the input protocol (top).

Motor control requires coordinated activation of many motor units. Evidences exist that this happens by subsequent triggering of muscle-synergies (*18, 19*), which control a subset of motor units into performing certain contraction patterns. Thus, multiple cell assemblies, embedded into the topography of the motor cortex, are involved in generating the correct activation sequence for the execution of a skill. Accordingly, when learning a new task, neural self-organization structures the network into the required assemblies. When practicing we check how far we have deviated from the desired goal and our nervous system derives from this an error signal used as feedback to guide the learning.

A similar feedback-controlled motor learning process can be shown in our network using a difficult task with a robot. Here we do not attempt to provide a detailed model of the human motor system. Rather we are concerned with the challenging problem to self-organize a network into functional units under feedback error control. For this we were choosing an accurate pick&place problem, which is very difficult to learn for small children as well as machines. The task was to insert a block into a very tightly fitting box (Fig. 3 A). To make this harder, we provide as reference signals for the learning only one single example action of putting the block into the box without having to rotate it and one other example of just rotating the block in right way and dropping it in the box without position change (see SOM). Creating a conjoint trajectory, thus, requires combining both these components. However, it can be observed that the robot fails by a substantial margin when it tries to do this prior to assembly growth (left panel in Fig. 3 C). Will training an assembly network lead to success? After all, it remains to be shown that there is no destructive cross interference occurring when several assemblies are active for generating a conjoint trajectory.

In order to solve this problem, we use two assemblies (Fig. 3 B), one for position and one for orientation and each computes one motor control function per degree of freedom used for trajectory generation (assembly *X*: three control functions for position, assembly *ϕ*: three for orientation (*20*)).

Two learning modes are possible. It is possible to learn both task components (position and orientation) simultaneously (not shown), or one can let the network learn alternatingly to put or rotate the object without performing both components together. We are here showing the second variant, because it demonstrates more clearly how the system behaves and how coexistent assemblies are formed.

Figure 3 D shows that the error signal drives the outgrowth of the assemblies, which gets slower as the error decreases until the system reaches the minimal cell assembly sizes required to successfully complete the task. Thus, such networks, in contrast to static structures, generate an optimal trade-off between performance and resources used. Remarkably, these cell assemblies are formed to coexist without leading to interfering activities and the final, total error is similar to the error of the two independent components (green shaded box in Fig. 3 D). This way, without learning this explicitly, a conjoint and accurate trajectory (*21*) is being executed by the robot on co-activation of both assemblies after only 8 learning trials (right panel in Fig. 3 C).

**Figure 3:**
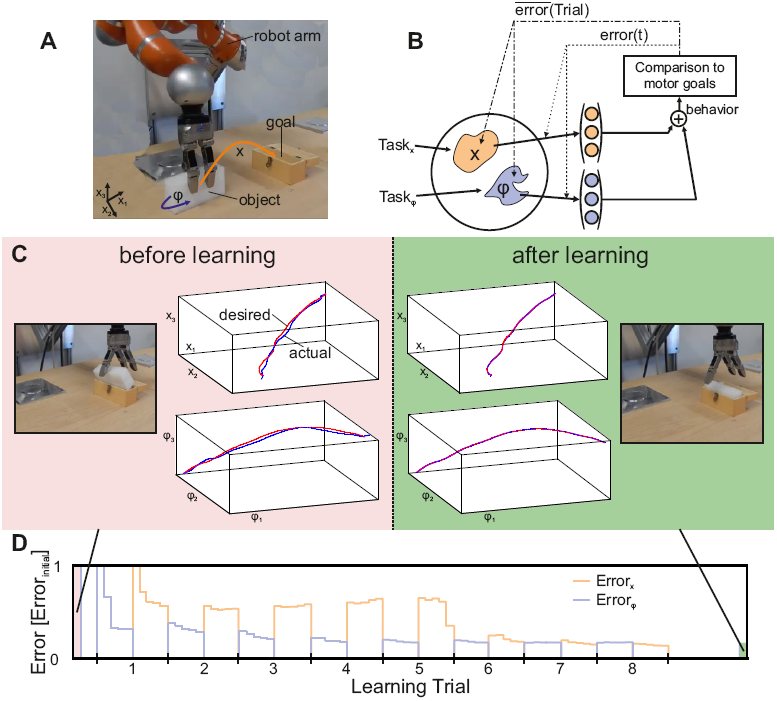
Highly accurate control of position and orientation in a robotic pick&place action is achieved by learning in coexistent, non-interfering assemblies. (A) Three degrees of freedom (DOF) each, for the trajectories for position *x* and orientation *ϕ*, need to be adjusted to an accuracy of about 1mm on each side to make the white block fit into the brown box by the robot. A human has provided two example trajectories by guiding the robot arm; one for position-only (DOFs: *x*_1_, *x*_2_, *x*_3_) and one for orientation-only (DOFs: *ϕ*_1_, *ϕ*_2_, *ϕ*_3_), which are used for reference. Their trajectories are encoded by dynamic movement primitives (DMPs (*22, 23*), see Materials and Methods), which use Gaussian kernels for every DMP equally spaced along the trajectory. (B) Learning needs to adjust the amplitude of the kernels until success. For this we grow and train two assemblies using average and detailed trajectory error (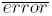), *error*(*t*)), respectively. Learning of position (input: *T ask_x_*) and orientation (*T ask_ϕ_*) is done independently and alternatingly. (C) Robot performance and trajectories before and after learning. (D) Both errors (for *x* and *ϕ*) drop into the success range with only eight learning trials. Please see SOM for protocol details and videos.

Previous works indicate that adaptation of the synaptic efficiency influences computational performance in neural networks (*10, 11, 17, 24, 25*), but the self-organization of a network into computationally powerful sub-structures poses a difficult and as yet unresolved problem.

In the current study efficient learning of a difficult motor skill was obtained by the combination of slow self-organization of the network into non-interfering cell assemblies with faster, error-driven acquisition of computational properties within these assemblies. As shown above, such systems are capable of combining substantial computational power with an economical use of network resources accommodating competition and/or coexistence of assemblies as determined by the inputs. This type of dynamic network restructuring is novel and not possible in static networks.

Structure, viz. cell-assembly-, formation must be adaptive but not volatile, which suggests that synaptic plasticity needs to be stabilized by processes that act on longer time scales – such as synaptic scaling. Our analytical results (see SOM) show that the combination of plasticity and scaling is indeed especially well suited to achieve coexistent assemblies and that it is difficult to achieve this by other plasticity rules (see SOM and (*9, 26*)).

Furthermore, it is known that synaptic plasticity and scaling act in many cortical areas (*27, 28*) and, thus, the interaction between cell assembly formation and their transient dynamics is not restricted to the here chosen example. For instance, experiments and models provide evidence for the existence of transient dynamics (*29*) within cell assemblies (*30*) in the prefrontal cortex, too.

An important additional aspect of such systems is that they will stay in a non-persistent activity regime without which computations will deteriorate (*31*). The transient activity present in our system is probably equivalent to the asynchronous irregular (AI) state in spiking networks (*15*), which, in the same way, provides rich and transient dynamics (*32*). This state is dominated by inhibition and, therefore, does not need any fine-tuning between inhibition and excitation (*24*); a strong property which is also shared by the here presented system. In particular, experimental data from delayed vibrotactile discrimination tasks are best described by a combined model of cell assemblies and transient dynamics (*33*). Thus, cell assemblies with transient neuronal dynamics seems to be ubiquitous feature in neural systems and the here presented results allow a better understanding of how such structures might be dynamically shaped into computationally powerful assembly networks.

## References and Notes

1. W. R. Ashby, An introduction to Cybernetics (Wiley, New York, 1956).

2. W. Maass, T. Natschläger, H. Markram, Neural Comput. 14, 2531 (2002).

3. H. Jaeger, H. Haas, Science 304, 78 (2004).

4. This paradigm is known as Reservoir Computing or Liquid State Machine (*2, 3*). Proofs exist that a rich-enough dynamic network of this kind can emulate a Turing machine and, thus, provide universal computational power (34).

5. J. J. Hopfield, Proc. Natl. Acad. Sci. USA 81, 3088 (1984).

6. G. Palm, A. Knoblauch, F. Hauser, A. Schüz, Biol. Cybern. 108, 559 (2014).

7. D. O. Hebb, The Organization of Behaviour (Wiley, New York, 1949).

8. G. Buzsaki, Neuron 68,362 (2010).

9. C. Tetzlaff, C. Kolodziejski, M. Timme, M. Tsodyks, F. Wörgötter, PLoS Comput. Biol. 9(10),e10003307 (2013).

10. S. Klampfl, W. Maass, J. Neurosci. 33(28),11515 (2013).

11. D. Sussillo, L. F. Abbott, Neuron 63,544 (2009).

12. R. C. Malenka, Nat. Rev. Neurosci. 4,923 (2003).

13. G. G. Turrigiano, K. R. Leslie, N. S. Desai, L. C. Rutherford, S. B. Nelson, Nature 391,892 (1998).

14. T. Keck, et al., Neuron 80,327 (2013).

15. N. Brunel, J. Physiol. 94,445 (2000).

16. K. D. Miller, Neuron 17,371 (1996).

17. H. Toutounji, G. Pipa, PLoS Comput. Biol. 10(3),e1003512 (2014).

18. M. C. Tresch, P. Saltiel, E. Bizzi, Nat. Neurosci. 2,162 (1999).

19. P. Saltiel, K. Wyler-Duda, A. D’Avella, M. C. Tresch, E. Bizzi, J. Neurophysiol. 85, 605 (2001).

20. Many more functions could be trained as reservoir network learning does only require large-enough networks for this (*2*). This way a more direct match to the framework of muscle synergies could be enforced, which is not of relevance here, though.

21. Note, the here-used trajectory generation framework (dynamic movement primitives (*22, 23*)) allows re-scaling and reshaping the learned trajectories for different configurations without having to re-learn from scratch. This is a long-known and important generalization property of such systems (*23*) and not in the core of our discussions. Other trajectory control methods can be used to the same ends, for example Gaussian Mixture Models (35).

22. J. A. Ijspeert, J. Nakanishi, S. Schaal, IEEE Int. Conf. Robot. (2002), pp. 1398–1403.

23. T. Kulvicius, M. Biehl, M. J. Aein, M. Tamosiunaite, F. Wörgötter, Robot. Auton. Syst. 61,1450 (2013).

24. G. Hennequin, T. P. Vogels, W. Gerstner, Neuron 82,1394 (2014).

25. C. Savin, J. Triesch, Front. Comp. Neurosci. 8,57 (2014).

26. C. Tetzlaff, C. Kolodziejski, M. Timme, F. Wörgötter, Front. Comput. Neurosci. 5,47 (2011).

27. D. E. Feldman, Annu. Rev. Neurosci. 32,33 (2009).

28. G. G. Turrigiano, Annu. Rev. Neurosci. 34,89 (2011).

29. V. Mante, D. Sussillo, K. V. Shenoy, W. T. Newsome, Nature 503,78 (2013).

30. K. Wimmer, D. Q. Nykamp, C. Constantinidis, A. Compte, Nat. Neurosci. 17(3),431 (2014).

31. This constraint is equivalent to the condition that the spectral radius of the network weights is smaller than one in echo-state networks (3).

32. S. Jahnke, R.-M. Memmersheim, M. Timme, Front. Comp. Neurosci. 3,13 (2009).

33. O. Barak, D. Sussillo, R. Romo, M. Tsodyks, L. F. Abbott, Prog. Neurobiol. 103,214 (2013).

34. W. Maass, Computability in context: computation and logic in the real world (London: Imperial College, 2010), chap. Liquid state machines: motivation, theory, and applications, pp. 275–296.

35. S. M. Khansari-Zadeh, A. Billard, IEEE T. Robot. 27,943 (2011).

36. The research leading to these results has received funding from the European Communitys Seventh Framework Programme FP7/2007-2013 (Programme and Theme: ICT-2011.2.1, Cognitive Systems and Robotics) under grant agreement no. 600578 as well as from the Germany Ministry of Science Grant to the Göttingen Bernstein Center for Computational Neuroscience, Project D1.

